# IPEV: Identification of Prokaryotic and Eukaryotic Virus-derived sequences in virome using deep learning

**DOI:** 10.1101/2023.05.28.541705

**Authors:** Hengchuang Yin, Shufang Wu, Jie Tan, Qian Guo, Mo Li, Xiaoqing Jiang, Huaiqiu Zhu

## Abstract

**Background:** The virome obtained through virus-like particle enrichment contain a mixture of prokaryotic and eukaryotic virus-derived fragments. Accurate identification and classification of these elements are crucial for understanding their roles and functions in microbial communities. However, the rapid mutation rates of viral genomes pose challenges in developing high-performance tools for classification, potentially limiting downstream analyses.

**Findings:** We present IPEV, a novel method that combines trinucleotide pair relative distance and frequency with a 2D convolutional neural network for distinguishing prokaryotic and eukaryotic viruses in viromes. Cross-validation assessments of IPEV demonstrate its state-of-the-art precision, significantly improving the F1-score by approximately 22% on an independent test set compared to existing methods when query viruses share less than 30% sequence similarity with known viruses. Furthermore, IPEV outperforms other methods in terms of accuracy on most real virome samples when using sequence alignments as annotations. Notably, IPEV reduces runtime by 50 times compared to existing methods under the same computing configuration. We utilized IPEV to reanalyze longitudinal samples and found that the gut virome exhibits a higher degree of temporal stability than previously observed in persistent personal viromes, providing novel insights into the resilience of the gut virome in individuals.

**Conclusions:** IPEV is a high-performance, user-friendly tool that assists biologists in identifying and classifying prokaryotic and eukaryotic viruses within viromes. The tool is available at https://github.com/basehc/IPEV.

## Introduction

Viruses or virus-like particles (VLPs) are the most abundant and diverse biological entities on earth, with an estimated 1031 particles, and even human feces contains 10^9^ VLPs each gram [1, 2]. Nextgeneration sequencing (NGS) technology has facilitated virome studies, leading to the discovery of new viruses and enhancing our understanding of the potential impact of viruses on environmental and human body microbiomes [3-6]. However, enriched sample approaches may result in the loss of host information about viruses, limiting downstream analyses. For instance, Wu et al. [7] discovered that ulcerative colitis disease states in patients are closely related to changes in the proportion of temperate phages in the gut virome. Such understanding relies on the fact that all eukaryotic viruses have been eliminated from the virome data. Moreover, the eukaryotic virome, as a critical component of the virome, likely plays a crucial role in host health and disease [8-12]. Recent research has revealed complex transkingdom interactions among eukaryotic viruses, bacteria, and the intestinal host [13-16]. To analyze these complex interactions between eukaryotic viruses and microbial communities, eukaryotic viruses need to be identified first. Thus, distinguishing prokaryotic and eukaryotic viruses from virome data is essential for a comprehensive understanding of the virus landscape [7, 13].

Nevertheless, the virome contains fast-mutating and diverse genetic elements, making it challenging to precisely identify prokaryotic and eukaryotic viruses from virome data. In addition, assembly tools may be limited due to mutation, recombination, and low or uneven sequencing coverage across the virus genome [17]. Numerous short and often insufficiently informative reads increase the difficulty of virus identification. Furthermore, the absence of a well-conserved genetic marker, such as the bacteria’s 16S rRNA gene, makes it difficult to construct a phylogenetic tree to distinguish between eukaryotic and prokaryotic viruses [18]. Virus identification typically involves aligning sequences against known viruses in genomic repositories, such as the National Center for Biotechnology Information (NCBI) Taxonomy Databases [19]. Despite the rapid increase in viral sequences, the number of viral reference sequences in public reference databases is still limited, constraining the effectiveness of sequence-based alignment methods. For instance, it has been estimated that there are millions of viral species, but the International Committee on Taxonomy of Viruses (ICTV) has currently recognized only 10,434 species to date [20].

Current computational tools for analyzing virome data focus on classifying the host and lifestyles of phages, such as VirSorter and HoPhage, which are designed to assign the host for a given phage contig using sequence similarity searching or supervised classification. Other tools, like DeePhage [7], have been developed to answer questions about the lifestyles of phages. While analyzing and classifying phages in virome data is a current focus, it is important to note that eukaryotic viruses also play a critical role in influencing host immunity and disease phenotype by infecting host cells and interacting with the bacterial microbiome through trans-kingdom interactions. However, existing methods for analyzing virome data are limited in distinguishing eukaryotic viruses from prokaryotic viromes, assigning hosts, and classifying them accurately [21]. Some methods, such as the Host Taxon Predictor (HTP) by Wojciech et al. [22], can be used to bridge the gap, differentiating between phages and eukaryotic viruses based on sequence information and nucleic acid type (such as DNA or RNA). Each virus sequence is represented by a vector of characteristics, including nucleic acid type, dinucleotide frequencies, relative dinucleotide, and trinucleotide frequencies. HTP predicts the taxonomy of viral hosts using four supervised machine learning techniques: k-nearest neighbors (KNN), support vector classifier (SVC), logistic regression (LR), and quadratic discriminant analysis (QDA). However, HTP’s performance is highly dependent on the nucleic acid type used, and some experimental protocols for virome research [23] may result in mixed datasets containing both DNA and RNA sequences, making it challenging to accurately determine the origin of these sequences. This uncertainty could potentially affect HTP’s performance in classifying viruses.

In this paper, we introduce IPEV, a high-performance, user-friendly tool for differentiating prokaryotic and eukaryotic virus from virome sequence fragments. We design a 64*64 sequence pattern matrix for numerical DNA sequences. IPEV employs a 2D convolutional neural network (CNN) to learn virus taxon information from this pattern matrix. Cross-validation tests demonstrate that IPEV significantly outperforms related methods in terms of F1-score metrics by approximately 20.8% while requiring only 1/50th the time of HTP in the same computing environment. We also designed a variety of homology layouts for independent sets, based on known sequence data, to assess IPEV’s generalization capabilities. Notably, IPEV outperforms HTP (KNN) by approximately 22% in terms of F1-score on an independent set with highly imbalanced labels when the sequence identity between the training and independent test sets is less than 30%. We applied IPEV to reanalyze longitudinal gut virome data from ten healthy individuals over 12 months, and it achieved the best performance in at least 90% of the samples compared to other methods. IPEV’s reanalysis revealed that the virome exhibits temporal stability beyond that observed in persistent personal viromes, thus enhancing our understanding of gut virome stability in individuals.

## Materials and Methods

### Dataset construction

In the absence of accurately host-annotated virome datasets that could serve as benchmarks, we generated simulated datasets based on reference genomes. Firstly, we downloaded the taxonomy ID list of viruses and corresponding host lineage from Virus-Host DB [24], and genome sequences from the NCBI database [25]. Our dataset comprised 11,022 eukaryotic virus and 5,051 prokaryotic virus genomes, including 113 archaeal viruses. We randomly divided 10,000 eukaryotic and 4,000 prokaryotic viruses for 5-fold cross-validation, while the remaining subset served as an independent test set for assessing generalizability. Considering the limitations of current mainstream sequencing technologies and the length constraints of assembled contigs, we simulated four contig length groups (A-D) using MetaSim (v0.9.1) [26] with “exact” pre-set and “Uniform” distribution type. The contig length groups were as follows: Group A (100-400 bp), Group B (400-800 bp), Group C (800-1,200 bp), and Group D (1,200-1,800 bp). We evaluated IPEV’s generalization ability using an independent test set consisting of 1,022 eukaryotic and 1,051 prokaryotic virus sequences. Using MetaSim, we generated 10,000 contigs and ensured low similarity against the training set using BLASTn (v2.7.1) following the above length groups. We total generated six low homology independent test sets (I1-I6) with varying query coverage and identity thresholds relative to the training set, as shown in Table S1.

We also evaluated the effect of sequencing errors on IPEV’s performance, and we generated a total of 10,000 contigs (1,200 to 1,800 bp) with 5%, 10%, and 15% sequencing errors based on the independent test set using MetaSim. To generate sequences with 5% base substitutions, the “Error Rate at Read Start” and “Error Rate at End of Read” from Metasim were both set to 0.05, while the “Insertion Error Rate” and “Deletion Error Rate” were both set to 0. The same settings were used to generate sequences with 10% base insertions or deletions, except that both the “Insertion Error Rate” and “Deletion Error Rate” were set to 0.5.

We analyzed longitudinal data from Shkoporov et al.’s [29] study to evaluate IPEV’s accuracy and the stability of gut virome data. We retrieved the raw human gut virome dataset from the NCBI SRA (accession number: PRJNA545408). This dataset included ten samples from healthy adults (subjects 916-925) collected over 12 months (T1-T12) through monthly synchronous samplings, with additional deeper sequencing of unamplified VLP nucleic acids at a one-time point (8th month) for each subject. We utilized the SPAdes (v3.13.0) [30] software to assemble short reads and conducted BLASTn searches against a bacterial database to eliminate bacterial contamination with an e-value of e-5, an identity of 50%, and query coverage of 90%. Our reference bacterial dataset comprised 20,003 complete prokaryotic genomes sourced from the NCBI RefSeq database, which consists of 19,629 bacterial genomes and 374 archaeal genomes. We aligned the genome sequences with reference virus sequences using BLASTn to assign virus taxon labels. The potential prokaryotic or eukaryotic viruses were inferred from viral contigs with an e-value of less than the cutoff 1e-4. Following Shkoporov’s personal persist virome (ppv) definition, we used cd-hit-est (v.4.8.1) with parameters c 0.8 aS 0.8 d 0 n 5 to cluster decontaminated contigs and defined clusters containing contigs from at least six months as ppv clusters. For subject 917, sampled for 11 months, we modified the ppv definition to contigs appearing in at least five months.

### Mathematical model of DNA sequences

In this study, we developed a sequence pattern matrix using the Sequence Graph Transform (SGT) model to extract meaningful information based on the relative positions of trinucleotides [31]. The pattern matrix functions as a numerical representation of frequency and order for trinucleotide pairs. We generated a trinucleotide set by combining three nucleotides to represent a DNA sequence (*S*) and calculated the weights of trinucleotides u and v using the following formula:

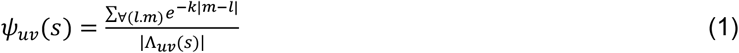

herein, *e*^−κ|*m*−*l*|^ represent the weight of a trinucleotide pair of *u* and *ν* at the position of *m* and *l*. The relative distances of a trinucleotide pair of *u* and *ν* are measured by |*m* −*l*|. |Λ _*uν*_| is the size of the set Λ _*uν*_. It represents the size of total (*u, ν*) pairs in the trinucleotide set from a DNA sequence. A schematic representation of the sequence pattern matrix can be found in Figure 1B. Finally, the DNA sequence is converted to a 64×64 matrix of relationship weights for the trinucleotide pair set.

**Figure 1.**
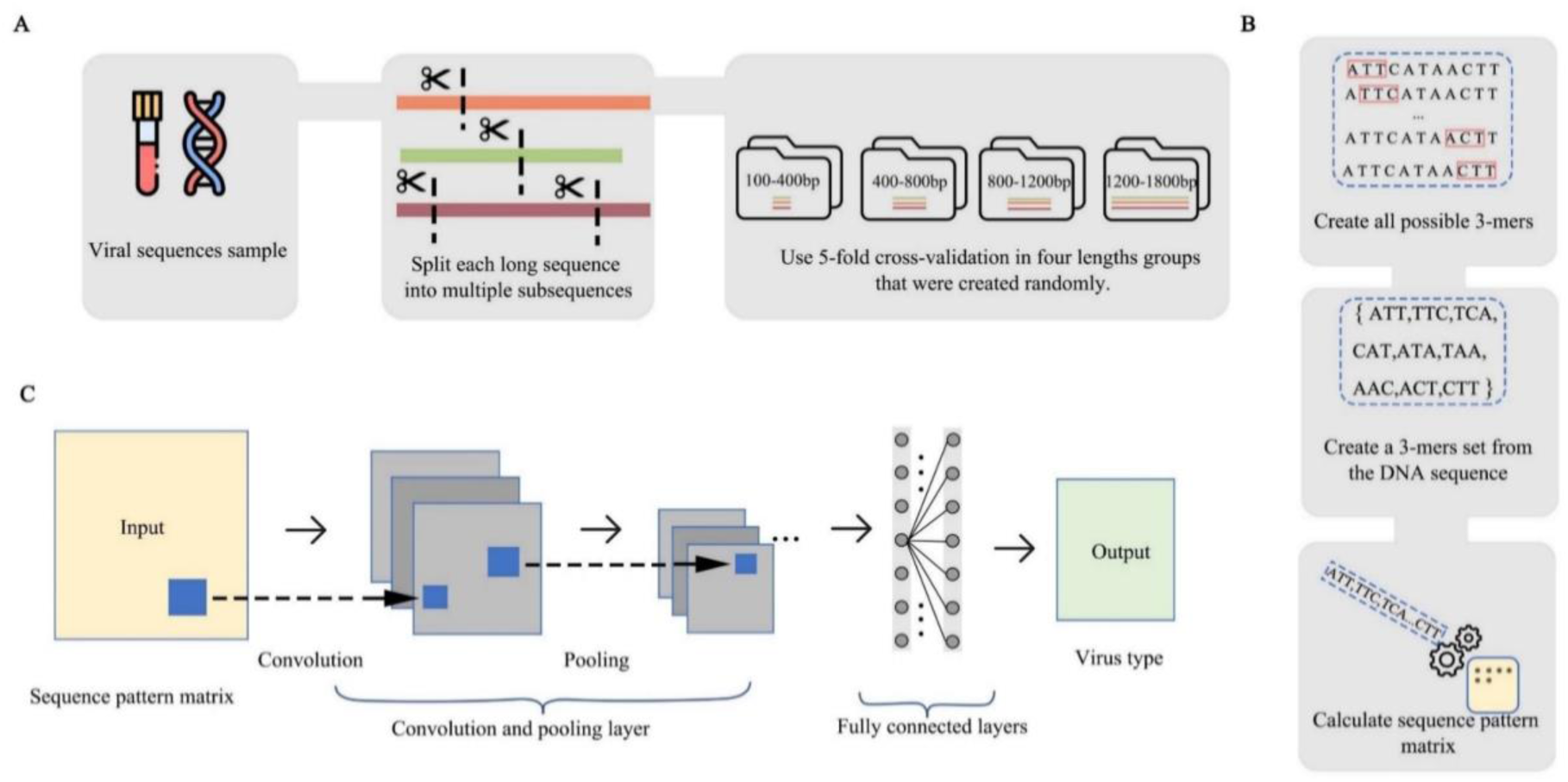
Workflow for extracting the sequence pattern matrix and using a deep learning neural network structure to predict taxon. **A**. The virus genomes are initially divided into five subsets, and then each subset is simulated to represent four groups with different contig lengths. **B**. Overlapping trinucleotides are used to represent the virus contigs. For example, if the nucleotides of the viral fragment are “ATTCATAACTT”, the trinucleotide set would consist of “ATT, TTC, TCA, CAT, ATA, TAA, AAC, ACT, CTT, CGT”. The trinucleotide set is then converted to a 64×64 sequence pattern matrix using a sequence pattern function. **C**. The IPEV tool employs a 2D CNN model as the classifier. The CNN model accepts the sequence pattern matrix as input and outputs a 1×2 array representing the likelihood of prokaryotic and eukaryotic viruses.

### Structure of deep learning neural network

We constructed a 2D convolutional neural network (CNN) to predict taxon information using sequence pattern matrices. The CNN has six layers: two convolution layers (with a 5×5 kernel and same padding), two max pooling layers (with a 2×2 filter), a fully-connected layer (which is flattened), and a fullyconnected layer followed by a softmax activation function. Here, the Conv2D layer takes sequence pattern matrix I of dimensions (L × *L*) as the sequence pattern matrix and generates total F feature maps as output by corresponding F (kernels) of dimensions *k*_1_ × *k*_2_ with the same padding. Those kernels were used to extract information on the viral sequence. Using ReLU (Rectified Linear Unit) as the activation function, the Conv2D layer output an F × *k*_1_ × *k*_2_ matrix *Y*^*c*^ and computes the f ^th^ feature map at the (*i*^th^, *j*^th^) location and the value is given as

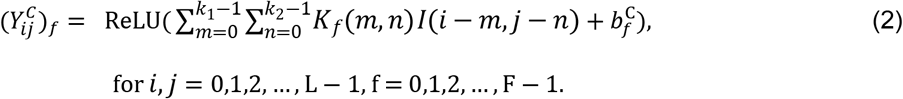

Where *k*^*f*^ and 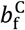 are a *k*_1_ × *k*_2_ weight matrix and a bias of the f^th^ kernel. Mainly, the ReLU function mentioned above is defined as follows:

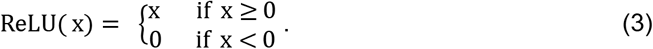

The next layer in the model is a Maxpooling layer, taking the maximum value over an input channel with a pooling size S1 × S1 and a stride size S2 × S2. The padding option is set to “valid”. The window is shifted along with each channel independently and can generate F new channels with the size of L′ × L′(L′ = ⌊(L − S1)/S2 + 1⌋). The Maxpooling layer outputs an L′ × L′ × F feature matrix *Y*^*M*^ and one of the pooling operations for a specific channel at the (*i*^th^, *j*^th^) location was calculated as

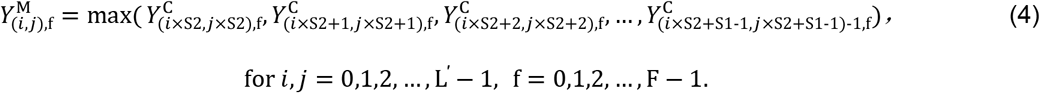

The features learned by a neural network in the Maxpooling layer are transferred to the Dropout layer. The output 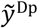 is formulated as

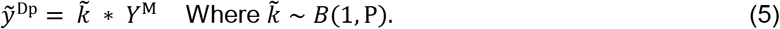

Here ∗ denotes an element-wise product. For any layer *Y*, The drop mask 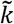 denotes an independent Bernoulli distribution with random variables, each having a probability *p* of 1. It could effectively reduce overfitting. We employed Flatten layer to convert all the elements ***Y***^Dp^ in the tensor into a *Y*^F^ one-dimensional array one by one. The Dense1 layer uses the ReLU function to output R units. It has an R×F weight matrix *w*^D1^ and an R-dimensional bias vector *b*^D1^. Each output unit is given as follows:

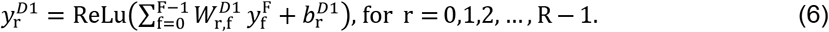

The Dense1 layer can generate an R-dimensional vector ***y***^D1^ While a Conv1D layer extracts features into different feature maps, and we use a SoftMax function as an activation function. The final layer is the Dense2 layer, which outputs only a 1 × 2 dimension array to represent the likelihood of phages and eukaryotic viruses. The output score is calculated as follows:

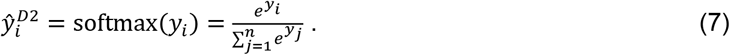

Moreover, the loss function is defined below

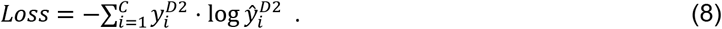

We employed the Adam optimizer (learning rate = 0.0001) and a batch size of 32 to train the neural network and update network weights (F = 64, M = 2, S1 = S2 = 2, P = 0.3, and R = 64). The architecture of the IPEV neural network is depicted in Figure 1C.

### Deriving viral taxon scores from subsequence predictions

To evaluate the performance of the neural network models, we carried out extensive training using 5-fold cross-validation within Groups A-D. When provided with a query virus fragment shorter than 1800 bp, the model from the corresponding length group was used. If the fragment length exceeded 1800 bp, it was first divided into 1800 bp windows, and the model from Group D was applied. The remaining subsequences, ranging from 0 to 400 bp, 400 to 800 bp, and 800 to 1,200 bp, were assigned to the first, second, or third models, respectively. The final prediction likelihood score was calculated as the weighted average of the scores obtained from each subsequence.

## Results

### Assessing the performance of ipev on viral genome fragments of different lengths using cross-validation

To assess the performance of IPEV and its comparative tools, we employed a 5-fold cross-validation procedure on Groups A-D. Considering that the HTP consists of four distinct classifiers, namely KNN, SVC, LR, and QDA, we compared their performances and selected KNN, which exhibited the best performance among the four classifiers, as our baseline for comparison. Our results demonstrate that IPEV outperforms HTP (KNN) in terms of average AUC value by 0.22, 0.20, 0.18, and 0.16, as illustrated in Figure 2. Additionally, we observed that the accuracy of the model’s predictions is directly proportional to the nucleotide sequence length. Therefore, the model’s ability to perform well on short genome lengths is a better indicator of the model’s reliability. IPEV showed superior performance, achieving an AUC value of 0.88 in Group A (100-400 bp), compared to HTP (KNN) with an AUC value of 0.66. Table 1 demonstrates that IPEV outperforms HTP in terms of mean accuracy by 17.1%, 19.5%, 19.9%, and 19.8% in Groups A-D. In binary classification, the sensitivity (recall) and precision measures evaluate the ability to predict positive and negative samples, respectively. HTP (KNN) tends to identify fragments as eukaryotic viruses in short fragments (100-400 bp) with relatively low sensitivity (precision = 65.0% ± 0.8% and sensitivity = 52.6% ± 0.5%). This suggests that HTP may miss some short phage fragments, which means that for some species with low abundance or insufficient sequencing depth, it may not be possible to assemble longer phage sequences, and HTP may fail to identify them, leading to false positives for eukaryotic viruses. Additionally, HTP (KNN) prefers to predict the virus’s host as prokaryotic in Group D (1,200-1,800 bp) with relatively high sensitivity (precision = 73.0 ± 1.4% and sensitivity = 80.8 ± 1.3%). IPEV, on the other hand, does not exhibit a specific preference. Notably, the F1 scores of IPEV in Group A-D outperform HTP (KNN), at 20.8%, 19.9%, 19.2%, and 18.6%. In conclusion, IPEV performs exceptionally well, with no apparent preference between eukaryotic and prokaryotic viruses in different length groups.

**Table 1.** The Average performance of IPEV and HTP (KNN) with 5-fold cross-validation for various sequence lengths (Percentage)

| Group | Tool | ACC | SN | SP | Precision | F1-score |
| --- | --- | --- | --- | --- | --- | --- |
| Group A | IPEV | 79.2±0.7 | 77.6±0.8 | 80.8±0.8 | 80.2±1.0 | 78.9±0.7 |
|  | HTP | 62.1±0.5 | 52.6±0.5 | 71.6±1.1 | 65.0±0.8 | 58.1±0.5 |
| Group B | IPEV | 88.8±0.6 | 88.7±0.3 | 89.0±1.3 | 89.0±1.1 | 88.9±0.5 |
|  | HTP | 69.3±0.7 | 68.0±1.0 | 70.5±1.7 | 70.0±1.1 | 69.0±0.5 |
| Group C | IPEV | 93.0±0.5 | 92.4±0.8 | 94.0±1.4 | 93.5±1.3 | 93.0±0.5 |
|  | HTP | 73.1±1.0 | 76.0±1.1 | 70.2±2.3 | 72.0±1.4 | 73.8±0.7 |
| Group D | IPEV | 95.2±0.5 | 95.0±0.6 | 95.3±1.0 | 95.4±0.9 | 95.2±0.5 |
|  | HTP | 75.4±0.9 | 80.8±1.3 | 70.1±0.2 | 73.0±1.4 | 76.6±0.6 |

\* Sensitivity = $TP / (TP + FN)$ , Specificity = $TN / (TN + FP)$ , Accuracy = $(TP + TN) / (TP + TN + FP + FN)$ , Precision = $TP / (TP + FP)$ , F1-score = $2 \times \text{Precision} \times \text{Recall} / (\text{Precision} + \text{Recall})$ , where TP, TN, FP, and FN respectively represent true positive, true negative, false positive, and false negative. As the method with the best performance in HTP, KNN is selected for comparison. The mean and standard deviation of 5-fold cross-validation are computed to elaborate on performance evaluation. Due to a lack of reconstruction between the train and validation set, the performance of HTP (KNN) is overestimated. (In this paper, prokaryotic viruses are treated as positive samples.)

**Table 2.** Comparison of IPEV and HTP (KNN) on the dataset-A (parameter: query coverage = 30%, identity = 30%).

| Group | Tool | ACC | SN | SP | Precision | F1-score |
| --- | --- | --- | --- | --- | --- | --- |
| Group A | IPEV | 0.7886 | 0.7622 | 0.7978 | 0.5673 | 0.6505 |
|  | HTP | 0.7261 | 0.5552 | 0.7855 | 0.4738 | 0.5112 |
| Group B | IPEV | 0.8602 | 0.8523 | 0.8629 | 0.6879 | 0.7613 |
|  | HTP | 0.7606 | 0.6822 | 0.7884 | 0.5332 | 0.5986 |
| Group C | IPEV | 0.9086 | 0.8863 | 0.9161 | 0.7778 | 0.8285 |
|  | HTP | 0.7929 | 0.7873 | 0.7948 | 0.5598 | 0.6543 |
| Group D | IPEV | 0.9449 | 0.9189 | 0.9533 | 0.8655 | 0.8914 |
|  | HTP | 0.8089 | 0.8176 | 0.8061 | 0.5793 | 0.6781 |

**Figure 2.**
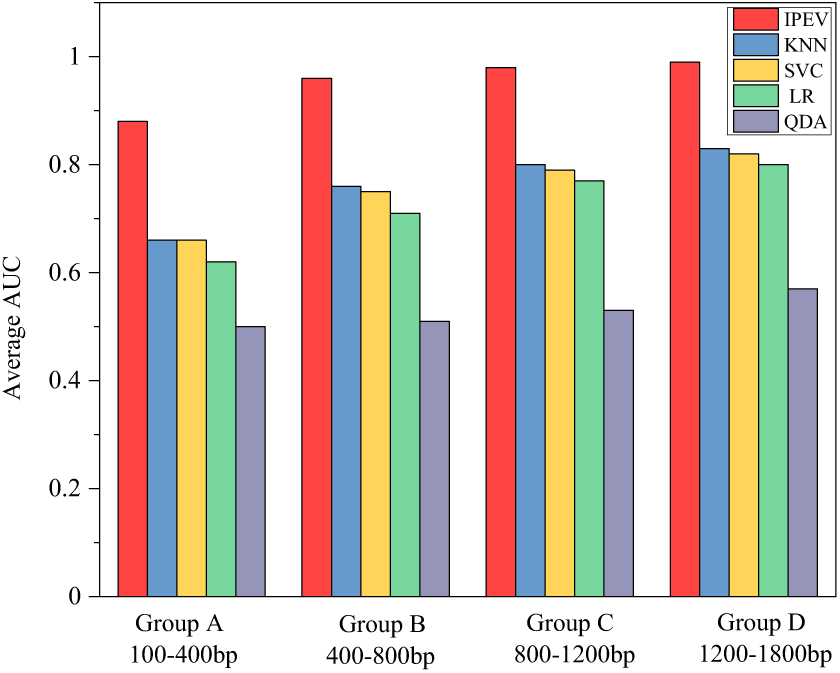
The average AUC value of IPEV and HTP (KNN, SVC, LR, QDA) for various lengths in k-fold (k=5) cross-validation. We plotted the training loss, validation loss, training accuracy and validation accuracy curves with respect to the number of epochs using 5-fold cross-validation. We observed that the model converged at 30 epochs, with training and validation losses and accuracies remaining consistent and overlapping. This indicates that the model achieved high performance while avoiding overfitting or underfitting on the data, as shown in Supplement Figure 1. To meet the high-throughput data requirements, we employed a parallel design for IPEV, which allows for more efficient processing of large datasets. Specifically, IPEV returns all taxon prediction scores in a single iteration with four loads of the neural network weight, significantly increasing the running speed. In contrast to HTP, where processing time is spent on file I/O operations, this approach enables IPEV to handle large datasets more effectively. To evaluate the model’s execution speed, we used the MetaSim software to simulate 10,000 sequence fragments. The results, shown in Figure S2, indicate that IPEV can reduce the time required by 50%. When using the same computational resources (CPU: Intel(R) Xeon(R), 20 cores, graphics cards name: NVIDIA Corporation GV100GL [Tesla V100 PCIe 32 GB).

### Evaluation of IPEV performance on novel viruses with low homology to known databases

The primary goal of taxon-predicting tools is to accurately predict newly discovered viruses, especially those with low homology to existing viral databases [22, 32, 33]. However, evaluating the performance of such tools is challenging due to the lack of accurate labels for new viral sequences. This study defines a novel virus as one with very low homology to known viruses. To assess the sensitivity of IPEV, we designed an independent test set with progressively increasing sequence identity to the IPEV training set. We describe the construction of this test set in the Materials and Methods section. Using a 30% identity and 30% coverage threshold, most high homology sequences were eliminated in an independent test set A. This highly labeled, unbalanced test set A contains Groups A, B, C, and D, which consist of 3,180, 3,152, 3,050, and 3,171 prokaryotic virus contigs and 1,106, 1,117, 1,011, and 1,036 eukaryotic virus contigs, respectively.

Despite removing sequences with high homology to the IPEV training set, IPEV still outperformed HTP. Specifically, in Group A (100-400 bp), IPEV reported a specificity of 0.80 and a sensitivity of 0.76. On the other hand, HTP (KNN) reported a sensitivity of 0.56 and a specificity of 0.79. These results suggest that compared to IPEV, HTP (KNN) tends to classify fragments as a eukaryotic virus when the sequence length is short. This may be because HTP (KNN) cannot obtain sufficient nucleotide frequency information from short sequences. As a result, the tool’s accuracy improves as the sequence length grows, resulting in a gradual decline in HTP’s preference. This conclusion is consistent with the results of a 5-fold cross-validation. In Group D, IPEV outperforms HTP (KNN) in specificity and sensitivity by 15% and 10%, respectively. As depicted in Figure 3, IPEV exceeded HTP (KNN) by 0.13, 0.11, 0.11, and 0.09 AUC values in Groups A, B, C, and D, respectively Furthermore, to assess the performance of IPEV, independent test sets B, C, D, E, and F were utilized. As the number of high homology sequences between the independent test set and the training set increases, the accuracy of IPEV is expected to improve. The results indicate that the similarity between the test and training sets is crucial in determining the classification performance. Table S6 demonstrates that in Group D of independent test set F, the highest accuracy that can be achieved with IPEV is 0.96, whereas the highest accuracy reported with HTP (KNN) is 0.80. It is important to note that the performance of HTP (KNN) is exaggerated. Additionally, the influence of imbalanced label data on the model’s performance is worth considering. Compared to HTP (KNN) in Groups A-D, the specificity and sensitivity of IPEV did not show significant changes as the number of imbalanced labels increased. These findings suggest that IPEV’s remarkably high prediction accuracy can identify potential virus taxon information even in an independent test set with low sequence homology to the training set. Importantly, our results demonstrate that IPEV does not exhibit bias in binary classification, as it maintains high performance even in the presence of imbalanced label data

**Figure 3.**
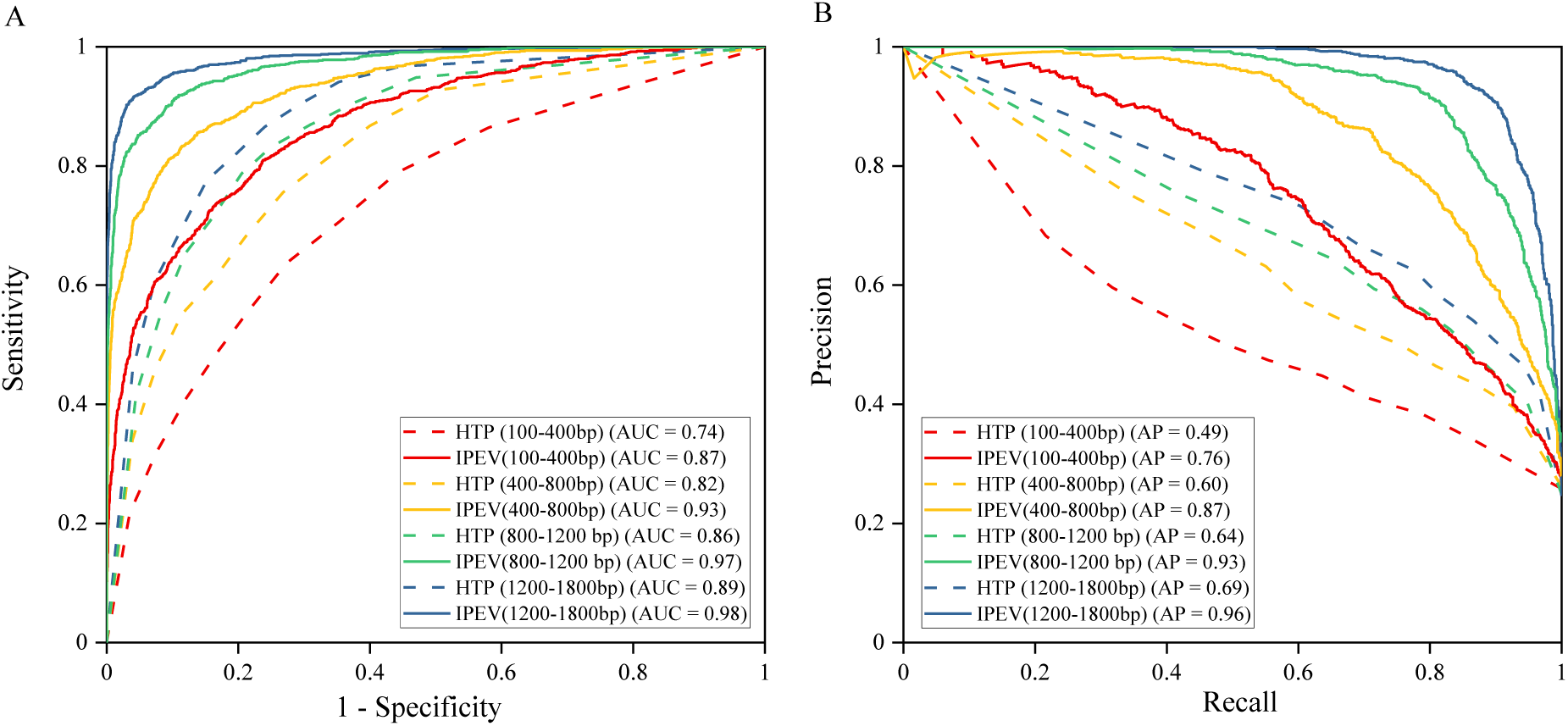
Performance comparison between IPEV and HTP (KNN) on dataset-A. **A**. The ROC (Receiver operating characteristic) curve demonstrates the discrimination capability, particularly in class-balanced test sets, with higher AUC values being preferred. B. The Precision-Recall curves serve as a measure of discrimination capability in class-imbalanced test sets, with AP representing the Average Precision.

### Evaluating IPEV performance on test sets with varying sequencing error rates

In this section, we evaluate the performance of IPEV and HTP on a dataset with varying levels of sequencing errors, including base insertion, deletion, and substitution. NGS errors can stem from factors like sequencing method and the experimental environment. Specifically, NGS has an error rate of 0.06% to 0.24% per base, while third-generation sequencing, such as PacBio, exhibit a relatively high error rate of 10% to 15% per base [34]. To evaluate the robustness of IPEV, we employed MetaSim to generate 2,000 short reads for both eukaryotic and prokaryotic viruses with different sequencing errors ranging from 1,200 to 1,800 bp. As shown in Table 3, the AUC values for both IPEV and HTP decrease with an increasing proportion of sequencing errors. When the substitution error rate reaches 15%, IPEV outperforms HTP (KNN) by approximately 16% in terms of AUC metric. This may be attributed to the sequence pattern matrix’s ability to tolerate errors. Furthermore, our results indicate that the percentage of sequencing errors introduced has only a slight effect on the IPEV’s performance, as the AUC value slightly decreases from 0.99 to 0.91 when the rate of error introduced by substitution increases from 0% to 15%. Nevertheless, IPEV remains the most accurate tool.

**Table 3.** comparison of IPEV and HTP performance on artificial datasets with varying error rates.

| Error Rate (%) | Base Substitute |  | Base Insert or Delete |  |
| --- | --- | --- | --- | --- |
|  | IPEV | HTP | IPEV | HTP |
| 0 | 0.99 | 0.82 | 0.99 | 0.82 |
| 5 | 0.98 | 0.80 | 0.98 | 0.80 |
| 10 | 0.95 | 0.77 | 0.97 | 0.80 |
| 15 | 0.91 | 0.75 | 0.95 | 0.78 |

### Evaluation of IPEV’s performance on functional protein sequences

In addition to evaluating short sequence fragments, we comprehensively evaluated the capability of the IPEV tool by incorporating protein sequences with functional annotations. One key aspect of understanding virus classification is identifying critical markers that play a role in the classification process, even though deep learning is often considered a black box. Given that receptor-binding proteins (RBPs) play a critical role in the adsorption of bacteriophages and the invasion of specific hosts, we hypothesized that they might contribute significantly to the prediction accuracy for phages. To test this hypothesis, we constructed a dataset consisting of 7,384 RBPs and corresponding negative samples, which were manually confirmed through Function, GO annotation, or Product containing RBP-related keywords, and additional filters. These protein sequences were obtained from various prokaryotic viruses, spanning seven orders and 28 families, such as Tubulavirales and Timlovirales.

The IPEV prediction results showed that RBPs significantly contributed to the prediction accuracy for phages. The predicted likelihood score by IPEV had a mean of 0.90 and a median of 0.98 for the RBP set, while for the non-RBP set, the mean was 0.77, and the median was 0.83, as shown in Figure 4.A (Wilcoxon rank-sum test, p-value < 2.2e-16). This finding suggests that the model can effectively learn host-related information to some extent, highlighting the importance of incorporating host-related information into phage prediction models. To further evaluate the performance of our model in predicting eukaryotic viruses, we specifically selected seven experimentally confirmed capping enzymes. Our analysis demonstrated that our model accurately predicted the likelihood of these protein sequences, with a score close to 1, providing evidence of its high accuracy and effectiveness in predicting eukaryotic viruses. Supplementary Table 2 presents further details of these results.

**Figure 4.**
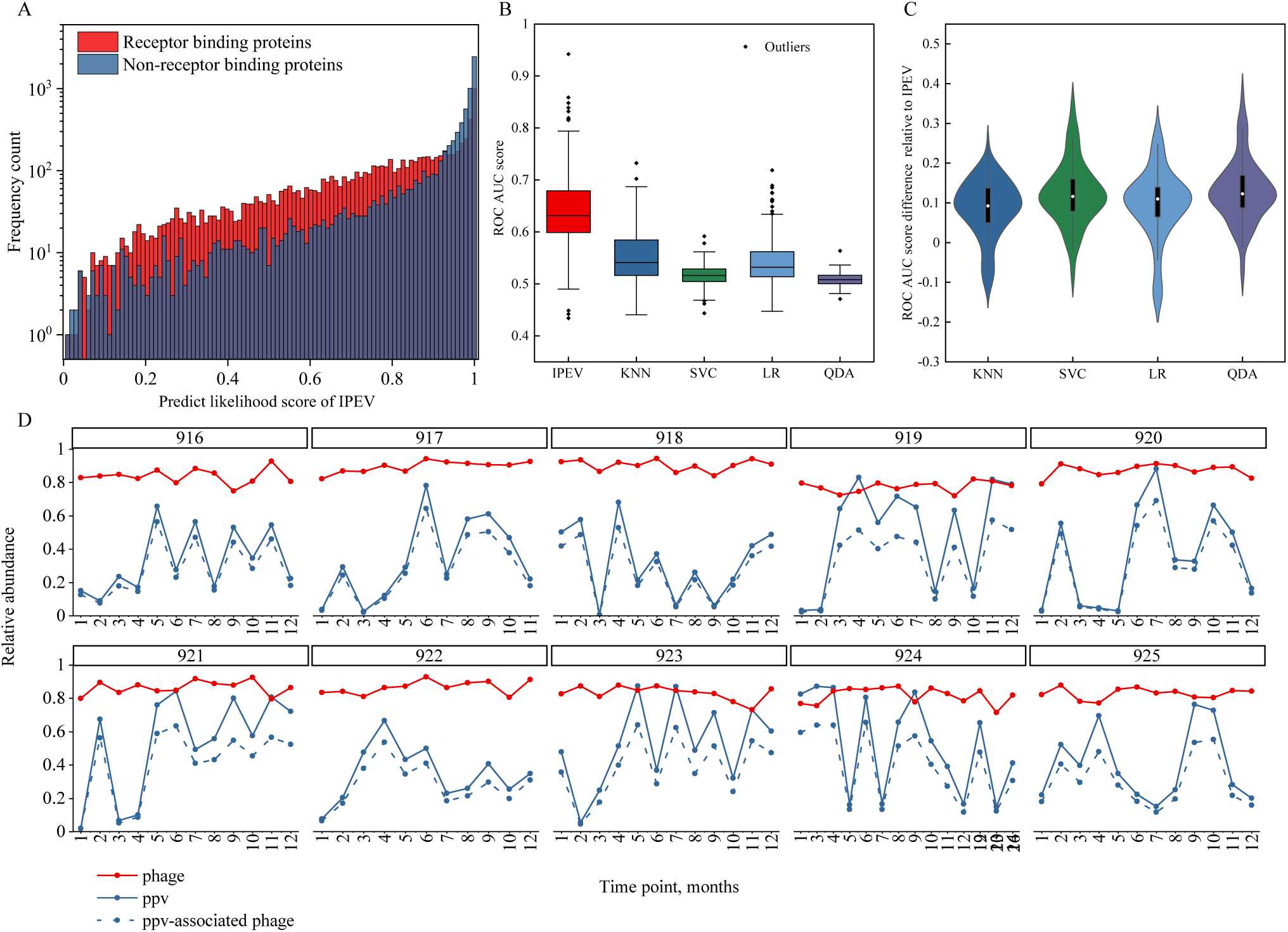
A. Histogram illustrating the predicted likelihood scores generated by IPEV for receptor-binding proteins (RBP) and non-RBP. B. Box plots representing the area under the curve (AUC) scores of the receiver operating characteristic (ROC) curves for IPEV, KNN, SVC, LR, and QDA. C. Violin plots displaying the AUC score differences of each tool relative in comparison to IPEV. D. Relative abundances of phages, PPV, and PPV-associated phages in the longitudinal data of subjects 916-925 as determined by IPEV.

### Applying IPEV to analyze the longitudinal gut virome in a cohort study

In this subsection, we present the results of our reanalysis, which includes an assessment of IPEV and related tool’s accuracy in the real virome and an exploration of the temporal stability of the healthy gut virome using longitudinal data over a year with the IPEV tool. The cohort comprised 130 samples from 10 subjects, and the raw data was processed and annotated as described in the Materials and Methods section. We employed IPEV and related tools for each sample to make predictions and compute AUC scores. Our analysis, as illustrated in Figure 4B, demonstrated that IPEV exhibited higher accuracy than HTP in more than 90% of the real virus samples and outperformed the other four models (KNN, SVC, LR, and QDA), with a mean AUC value of 0.64, which was significantly higher than the mean AUC values of 0.55, 0.51, 0.54, and 0.51 for KNN, SVC, LR, and QDA, respectively. Additionally, the median AUC for IPEV was 0.63, which was significantly higher than the median AUC values of 0.54, 0.52, 0.53, and 0.51 for KNN, SVC, LR, and QDA, respectively. The median difference between the AUC of IPEV and KNN, SVC, LR, and QDA for each sample is shown in Figure 4C, with median differences of 0.09, 0.12, 0.11, and 0.12, respectively. It is worth noting that QDA is equivalent to random guessing. Although the results on the simulated dataset were not as impressive as those on the real virome data, this discrepancy could potentially be attributed to the fact that the assembled sequences in our 130 samples predominantly consisted of short sequences, with sequences below 500bp accounting for 80% of the total. Despite this challenge, our model still exhibited the highest accuracy in identifying viruses within the real virome, showcasing its effectiveness and robustness. The human gut virome, characterized by its immense diversity and abundance of virus particles, is often referred to as the “dark matter” of the gut microbiome. The gut feces contains up to approximately 10^9^ VLPs per gram, yet only a fraction of the virus genomes, ranging from 14.2% to 56.6%, can be annotated. Despite its limitations, previous research has successfully identified a highly individualized and persistent fraction in the gut virome, known as the personal persistent virome (PPV). Additionally, this research has observed temporal stability in the virus components at the individual level. To further explore the virome, we employed IPEV to ab initio annotate contigs in the virome while excluding bacterial contamination. The results showed that the average coefficient of variation of the phage virome phage was significantly lower (0.04 ± 0.01) compared to the PPV (0.58 ± 0.05), indicating a high degree of temporal stability of phage, as illustrated in Figure 4C. Our findings are consistent with prior studies and provide additional support at a higher taxonomic level for the hypothesis that a “kill-the-winner” mechanism may be at work in in phage strains and substrains particularly. This mechanism prevents any one species from dominating and allows numerous species to coexist, so improving the diversity of the gut ecosystem and boosting its general resilience [35]. In contrast, the PPV is dominated by lytic life cycles and composed of virulent crAss-like and Microviridae phages that infect major representatives of the bacterial microbiota, resulting in higher variations than those seen in gut ecology. We also observed limited transient disturbance in the phage virome component in subjects 916 (T1 and T7) and 922 (T3, T5, and T8) with antibiotic usage. Interestingly, despite differences in individual gender, BMI, and lifestyle factors such as smoking and alcohol consumption, there were similar relative abundances of phage virome among individuals. In contrast, PPV had highly individualized relative abundances among individuals. Furthermore, we used IPEV to calculate the contribution of phage and eukaryotic viromes to PPV variability, demonstrating that phages are the primary component and dominate PPV fluctuations. Our results further enhance our understanding of the stability of the gut virome.

### Discussion and Conclusion

As mobile genetic elements (MGEs), viruses can transfer genes between different cells or species through horizontal gene transfer (HGT), thereby influencing the diversity and complexity of viral genes. The high level of viral diversity presents a significant challenge for virus identification tools. In this study, we introduce IPEV, a novel method that utilizes a sequence pattern matrix and a 2D convolutional neural network to distinguish prokaryotic and eukaryotic viruses-derived sequence fragments. To the author’s best knowledge, IPEV is the first de novo identification algorithm tool developed to addres this type of problem for virome data.

IPEV offers several advantages over traditional genomics techniques, such as k-mer methods and one-hot encoding. By integrating position and frequency information of 3-mers into a sequence pattern matrix, IPEV not only enhances the efficiency of the neural network model but also preserves valuable information about the order and position of trinucleotide. Moreover, we parallelize the sequence processing to enhance the speed. Our results demonstrate that IPEV can reduce analysis time by 50-fold, taking only 9.6 minutes to analyze 20,000 sequences of length 1,200-1,800. This improvement inefficiency makes IPEV a valuable tool for high-throughput data exploration in genomics research.

To ensure the generalization evaluation of our tool, we opted not to use the traditional approach of dividing the dataset into training and testing sets based on the date the sequence was discovered [36-38]. This method can lead to the inclusion of sequences with high homology to the training set in the test set, inflating the algorithm’s accuracy and making it challenging to accurately assess its performance. Instead, we used a series of thresholds to gradually remove sequences from the test set that are homologous to the training set. while artificial contigs with less than 30% sequence homology against the training set, IPEV achieved an average AUC of 0.98, indicating that our model learned valuable information and did not solely rely on sequence similarity for prediction. Our results suggest that IPEV can generalize well and provide reliable predictions for prokaryotic and eukaryotic viruses-derived sequence fragments.

We aimed to interpret the neural network based on its probability scores for phage prediction. The RBP is essential for the specific interaction of the bacteriophage with the host in phage prediction. Thus, we selected all RBP genes annotated with ‘Fiber’ and ‘Spike’, and an equal number of non-RBP genes, from two datasets, IPEV and another dataset. The score distribution of IPEV differed between the two datasets, with IPEV showing a higher score distribution in the former dataset. This indicates that RBPs with host information have more influence on phage prediction in IPEV. As the neural network model learns from the well-annotated data, IPEV can also discover some information about the virus host. This can help us understand neural network better in related task.

We employed IPEV to thoroughly analyze longitudinal data of the gut virome, which is comprised of both eukaryotic and prokaryotic viruses. These viruses infect human cells, microorganisms (such as bacteria, fungi, and archaea), and plant viruses that mainly originate from the environment and diet.Despite the limited annotations of these viruses, we successfully annotated all the sequences and found that the phage community is resilient to environmental perturbations, and the impact of the “kill the winner” theory on the phage community is at a low taxonomic level. This finding is consistent with previous studies. Our study supports the hypothesis that the “kill the winner” theory promotes a more diverse species community, which enhances ecosystem resilience. Furthermore, we observed that changes in ppv were primarily driven by phage, particularly those that target representative gut microbiota, resulting in significant fluctuations in ppv abundance. Our tool has a limitation in that it does not perform downstream virus classification or specific virus-host prediction for virome data. To fully comprehend the impact of viruses on microbial communities and hosts, a more specific and systematic classification is necessary. We look forward to the development of more tools that can reduce the degree of virome dark matter and improve our understanding of the virome.

## Availability of data and materials

Our study contains only publicly available viral completed genome sequence, reference bacterial completed genome sequence, IPEV program, and a detailed tutorial that can be publicly available at GitHub [https://github.com/basehc/IPEV] or Zhu Lab home page[http://cqb.pku.edu.cn/ZhuLab/IPEV] under GNU General Public License V3. Code and data for transparent and reproducible results were documented on GitHub [https://github.com/basehc/IPEV_analysis].

## Author contributions statements

H.Q.Z., H.C.Y., and S.F.W. conceived and designed the project; H.C.Y., J.T., and S.F.W. constructed the datasets; H.C.Y. and S.F.W. wrote and optimized the model of IPEV; H.C.Y. and S.F.W. performed the data analysis and design of the pipeline and prepared all the figures and tables; H.C.Y. drafted the manuscript. H.C.Y., H.Q.Z., S.F.W., M.L., X.Q.J, and J.T. revised and edited the manuscript, and all authors proofread and improved the manuscript.

## Acknowledgments

This research was supported through computational resources provided by the High-Performance Computing Platform of the Center for Life Science of Peking University. This work was supported by the National Key Research and Development Program of China (2017YFC1200205) and the National Natural Science Foundation of China (32070667, 31671366).

